# Ongoing brain rhythms shape I-wave properties in a computational model

**DOI:** 10.1101/205450

**Authors:** Natalie Schaworonkow, Jochen Triesch

## Abstract

**Background:** Responses to transcranial magnetic stimulation (TMS) are notoriously variable. Previous studies have observed a dependence of TMS-induced responses on ongoing brain activity, for instance sensorimotor rhythms. This suggests an opportunity for the development of more effective stimulation protocols through closed-loop TMS-EEG. However, it is not yet clear how features of ongoing activity affect the responses of cortical circuits to TMS.

**Objective/Hypothesis:** Here we investigate the dependence of TMS-responses on power and phase of ongoing oscillatory activity in a computational model of TMS-induced I-waves.

**Methods:** The model comprises populations of cortical layer 2/3 (L2/3) neurons and a population of cortical layer 5 (L5) neurons and generates I-waves in response to TMS. Oscillatory input to the L2/3 neurons induces rhythmic fluctuations in activity of L5 neurons. TMS pulses are simulated at different phases and amplitudes of the ongoing rhythm.

**Results:** The model shows a robust dependence of I-wave properties on phase and power of ongoing rhythms, with the strongest response occurring for TMS at maximal L5 depolarization. The amount of phase-modulation depends on stimulation intensity, with stronger modulation for lower intensity.

**Conclusion:** The model predicts that responses to TMS are highly variable for low stimulation intensities if ongoing brain rhythms are not taken into account. Closed-loop TMS-EEG holds promise for obtaining more reliable TMS effects.

## Introduction

Transcranial magnetic stimulation (TMS) is a non-invasive technique for stimulating cortical circuits through the use of externally applied magnetic fields. Currently, its therapeutic potential is limited, as the effects of a given stimulation protocol cannot be reliably predicted. TMS effects are notoriously inconsistent, with the same stimulation protocol inducing plasticity effects in opposite directions. This holds not only across subjects but even within a single subject [1, 2, 3, 4]. A major goal of current research is therefore to better understand and eliminate the sources of this variability. While the spatial accuracy of TMS has markedly increased through the use of neuronavigation techniques [5], an optimization of temporal aspects of stimulation could yield more consistent plasticity effects.

In this context, several recent TMS-EEG studies have shown a dependence of TMS responses on features of brain activity at stimulation onset in post-hoc evaluation. This was shown for slow oscillation up- and down-states in sleep [6], oscillatory power [7, 8, 9] and phase of different oscillatory rhythms [10, 11, 12], see [13] for review. The recent development of real-time triggered TMS-EEG systems now allows to precisely control stimulation based on measurements of ongoing brain activity [14]. Therefore, an improved understanding of the interaction between ongoing brain activity and TMS on cortical circuits is called for to support the development of more reliable plasticity induction protocols based on real-time triggered TMS-EEG.

When TMS is performed over the motor cortex, two main read-outs can be used to quantify the effects. First, TMS induces a twitch of the targeted muscle, which causes a motor evoked potential (MEP), measurable by electromyography. Second, in patients with electrodes implanted in the epidural space [15] a characteristic high-frequency discharge can be recorded following TMS, termed D- & I-waves. While such recordings are rare, understanding I-wave dynamics gives insight into how cortical circuits are driven by TMS. Here we use computational modeling to better understand how ongoing rhythmic brain activity may influence the effects of TMS on cortical circuits. We extend a previous model of TMS-induced D- & I-waves [16] to investigate the joint effect of background activity and stimulation pulses on cortical responses.

In the extended model, we test for systematic effects of ongoing rhythmic brain activity, such as phase and power of the *μ*-rhythm, on different measures of the TMS-induced responses. We find that the effects of TMS are modulated by phase and power of the oscillatory background activity at the time of stimulation and that TMS produces the largest responses when triggered at times when pyramidal neurons are highly depolarized. Furthermore, we show that higher response variability is observed for higher oscillatory power and discuss the implications of these findings for TMS-EEG. Additionally, we show that phase and power affect I-wave amplitudes and raw spike counts to a different degree, which reveals limitations of using compound signals to evaluate TMS effects.

## Material and Methods

### Model assumptions

The model generates D- and I-waves through an interaction between membrane mechanisms and dendritic processing of synaptic inputs. It makes two crucial assumptions: first, I-wave periodicity depends on the length of the refractory period of corticospinal tract neurons in layer 5 (L5) of motor cortex. The refractory period determines the maximum firing rate these neurons can achieve. Single-unit recordings during direct motor cortex stimulation [17] show that neurons involved in I-wave generation are capable of firing with a frequency of ~670 Hz, corresponding to a refractory time of 1.5 ms, as also supported by more recent studies [18]. The second assumption is that the arrival of synaptic input to these neurons is distributed across their dendritic trees. In combination, these properties result in high-frequency discharge patterns after TMS.

### Model structure

The circuit model consists of layer 5 (L5) neurons each receiving input from 300 layer 2/3 (L2/3) neurons, as illustrated in Fig. 1 (left). The L5 neurons have a detailed morphology (reconstructed by [19]), consisting of 188 compartments. The L2/3 neurons are modeled as single cylinders, the diameter equal to the length (taken from [20]). They are of excitatory and inhibitory type, with ratio 4:1. The L2/3 neurons project with one synapse each onto the dendritic branches of the L5 neuron in a feed-forward manner. For simplicity, we assume that the synapses are distributed uniformly across the dendritic tree [21], with a constant density per unit length. The conduction delay between L2/3 and L5 neurons is set to 1 ms. This will determine the latency between the D- and the first I-wave. [22] investigated this delay with single-unit recordings and found a symmetric distribution of latencies around 1 ms. The full model comprises 100 L5 neurons with a myelinated axon, consisting of nodes of Ranvier and myelin sheaths. Model output is evaluated with two different measures, TMS-induced spikes and TMS-induced I-wave amplitudes. The simulated epidural recording of I-waves is based on the mean membrane potential of all L5 neurons at the middle of the axon. The mean membrane potential is convolved with a Gaussian kernel (σ =0.25 ms) to account for different L5 conduction delays. I-wave peak amplitudes were calculated as peak value minus next trough value [23]. Peaks and troughs are identified with a local extrema search, which determines the peak maximum/minimum amplitude value by comparing each data point in the time series with its neighbors in a window of 0.5 ms. If it is larger/smaller than all neighbors, it is identified as a maximum/minimum. This reliably determines I-wave peaks and subsequent troughs.

**Figure 1:**
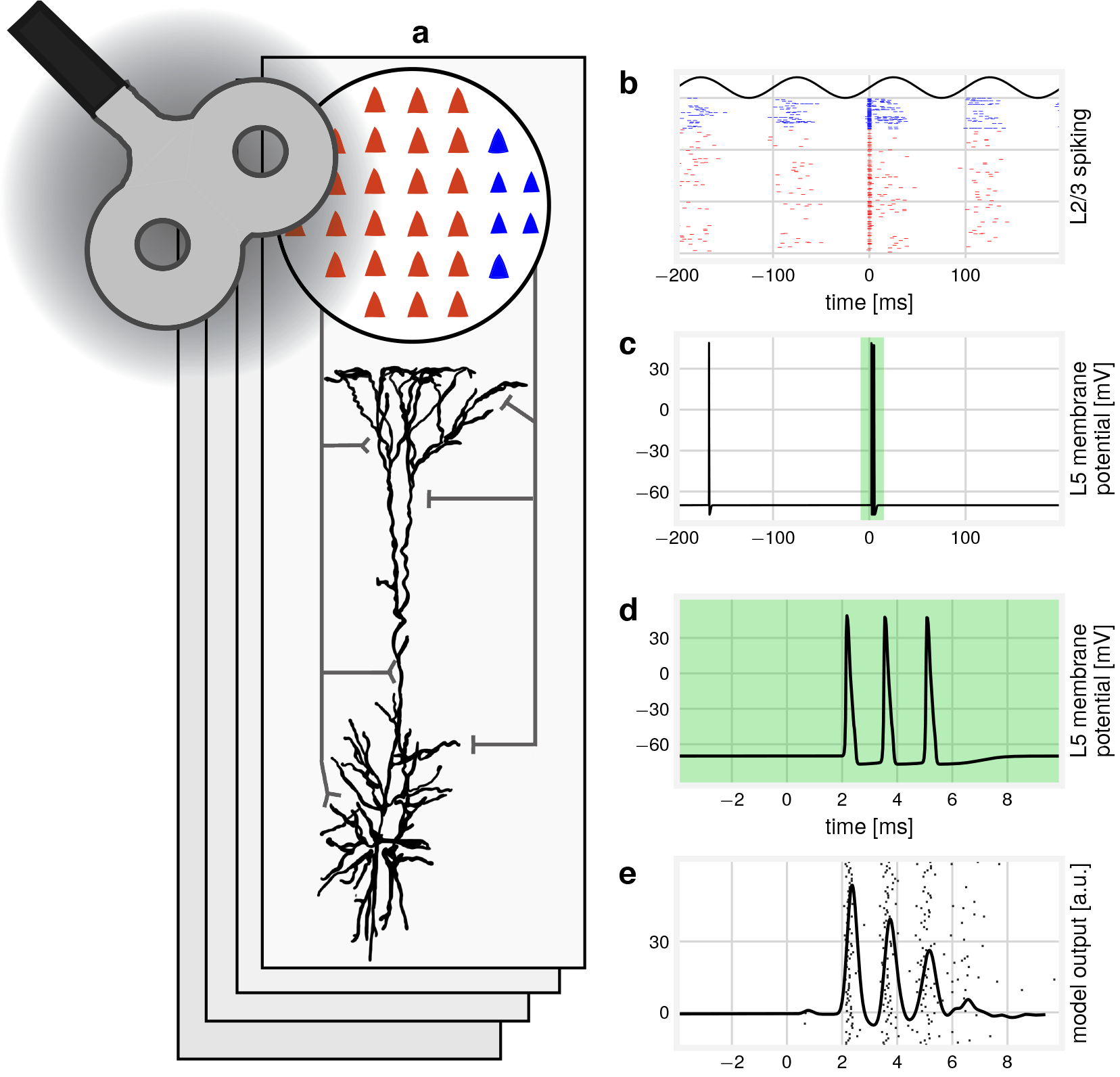
Network structure and activity. (a) Model structure: 60 inhibitory and 240 excitatory L2/3 neurons connect to a multi-compartmental L5 neuron in a feed-forward manner. 100 of these subnetworks are combined to produce the model output. (b) Example L2/3 firing patterns over time. The inhomogeneous Poisson input is modulated by a 10 Hz sine wave. At time=0, a TMS pulse is applied. (c) The resulting L5 firing, recorded in the axon. The green shaded area marks the response to the TMS pulse. (d) Spiking behavior around the time of the TMS pulse. (e) Example D- and I-wave model output, pooled over all subnetworks. In the background, the generated spikes are displayed. The dispersion of spike times increases for subsequent I-waves, resulting in a smaller amplitude for *I*_2_- and *I_3_*-waves.

### Model dynamics

L5 membrane potential dynamics are governed by Hodgkin-Huxley type equations in all compartments. L2/3 current types consist of sodium and delayed rectifier currents for action potential generation and a slow voltage-dependent potassium current for spike-frequency adaptation. Current types are specified in Table 1. Synaptic conductance changes are implemented as a sum of exponentials, allowing for independent rise and decay times. Four different types of synapse dynamics are used: fast and slow excitation (AMPA- and NMDA-type dynamics) and fast and slow inhibition (GABA_A_ and GABA_B_-type dynamics). These are reflected in different rise and decay times for the respective synapses. Time constants, reversal potentials and conductances are given in Table 1. Conductances are multiplied by synaptic weights drawn from a lognormal distribution to model the lognormal-like distribution of synaptic efficacies in the cortex [24]. To enable rhythmic spontaneous firing, peak conductance and reversal potential for GABA_B_ was adjusted, as well as the peak conductance for AMPA (see model summary table), as the original values [16] resulted in inhibition outweighing excitation, with no spontaneous firing possible. The L2/3 to L5 synapses exhibit a form of short-term synaptic depression. After each presynaptic spike the maximal synaptic conductance is decreased by a depression factor *d* = 0.5, with an exponential recovery with time constant *τ*_D_ = 200 ms [25].

### Spontaneous activity and stimulation

To model rhythmic spontaneous activity, L2/3 neurons receive input from an inhomogeneous Poisson process, modulated by a 10 Hz sine wave, representing rhythmic thalamic input [26, 27]. We model three different levels of oscillation power. Example input spikes to L2/3 neurons are illustrated in Fig. 2. The oscillatory drive leads to rhythmic firing in the L2/3 neurons. The inhibitory L2/3 firing rates are about three times higher than excitatory rates [28], see Fig. 1 (right) for an example of baseline firing. Spontaneous firing rates for L2/3 neurons are 2.0 ± 1.6 Hz for excitatory and 8.4 ± 4.7 Hz for inhibitory neurons [29]. The rhythmic firing in the L2/3 neurons results in subthreshold oscillations of the somatic L5 neurons’ membrane potentials [30]. The individual mean firing rate of the L5 neurons is 1.8 ± 2.3 Hz, meaning that individual neurons do not fire in each cycle of the oscillation. To investigate any state-dependent effects of TMS, we vary the timing of the TMS pulse relative to the phase of the background oscillation. We also vary oscillatory power and TMS intensity.

**Figure 2:**
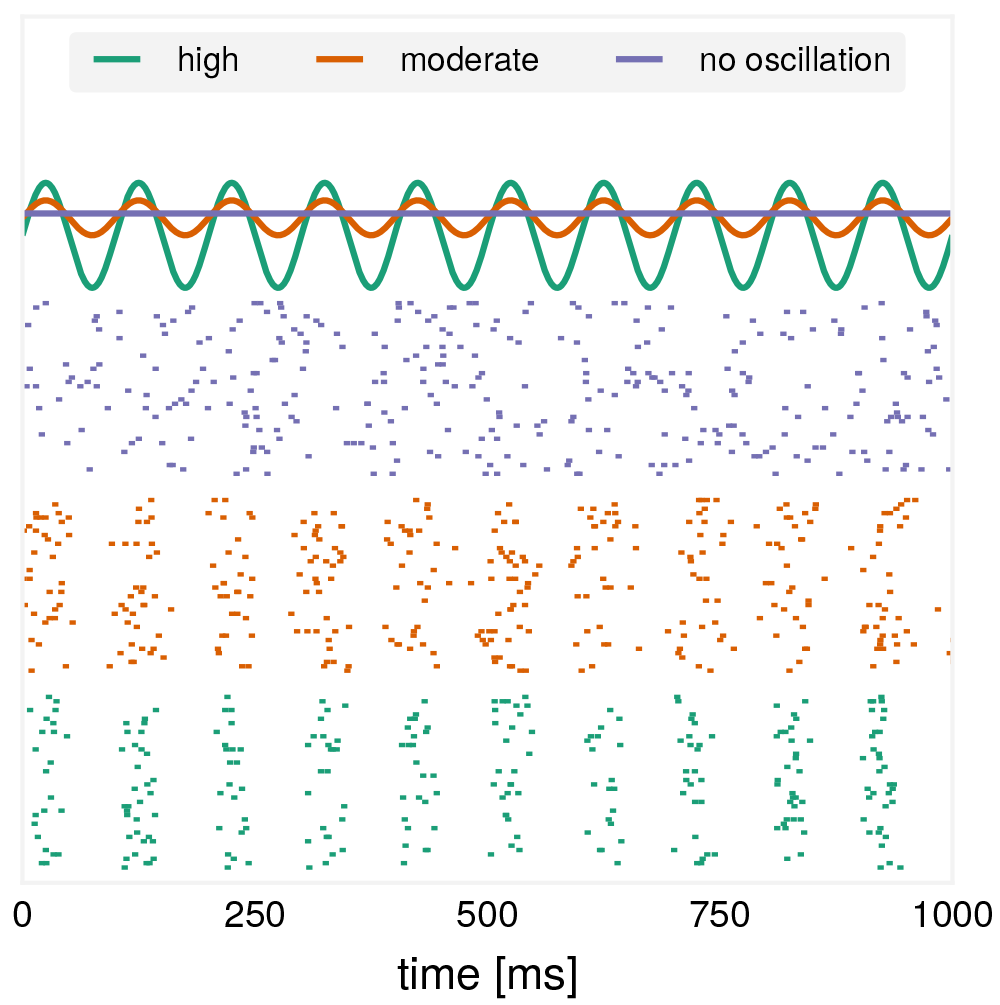
Network input: L2/3 neurons receive rhythmic input with varying power of the oscillatory rhythm. Top: The stimulus generating function for three different conditions: constant firing (purple), firing with an oscillation of moderate (orange) or high power (green). The power was varied by changing the offset of the generating sine, while maintaining the same mean firing rate for each condition by varying the amplitude. Bottom: Example raster plots for each condition.

Due to neuronal membrane time constants and conduction delays, membrane oscillations of L5 neurons are delayed with respect to the oscillatory driving input to the L2/3 neurons. We estimate this delay by fitting a sine function to the spontaneous membrane potential oscillation of the L5 cells. The L5 membrane oscillations are delayed by 14.0 ms with respect to the driving oscillation. All results are plotted against the phase of the subthreshold oscillations in the somatic membrane potential of the L5 neurons (subsequently referred to as: L5 phase). Because the membrane is most depolarized at L5 phase 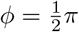, also the probability of firing is highest at this phase. Conversely, the probability of firing activity is lowest at L5 phase 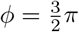.

TMS is modeled as a current injection into the somata of the L2/3 and axon of L5 neurons. The current has a temporal structure, modeling the output of an underdamped RLC circuit of commercial TMS systems. To model different excitability levels of neurons, depending on their location and orientation with respect to the coil [31], the maximal amplitude of the pulse *I*_max_ is drawn from an exponential distribution with scale parameter *β*. To model different stimulation intensities, *I*_max_ is multiplied by gain factor γ. The parameter *β* is a free parameter, adapted to the firing threshold for each neuron type to cover a broad range from no to a high number of neurons activated by TMS. This results in three parameters: *β*_*L*5_, 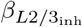 and 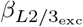. We make the assumption that inhibitory L2/3 neurons are easier to activate by TMS than excitatory L2/3 neurons [32]. For the L5 neurons the fraction of activated neurons is even smaller, reflecting the fact that a higher stimulation intensity is needed to evoke a D-wave compared to I-waves. The fraction of activated neurons for each type is illustrated in Fig. 3. The stimulation intensity is increased by increasing γ.

**Figure 3:**
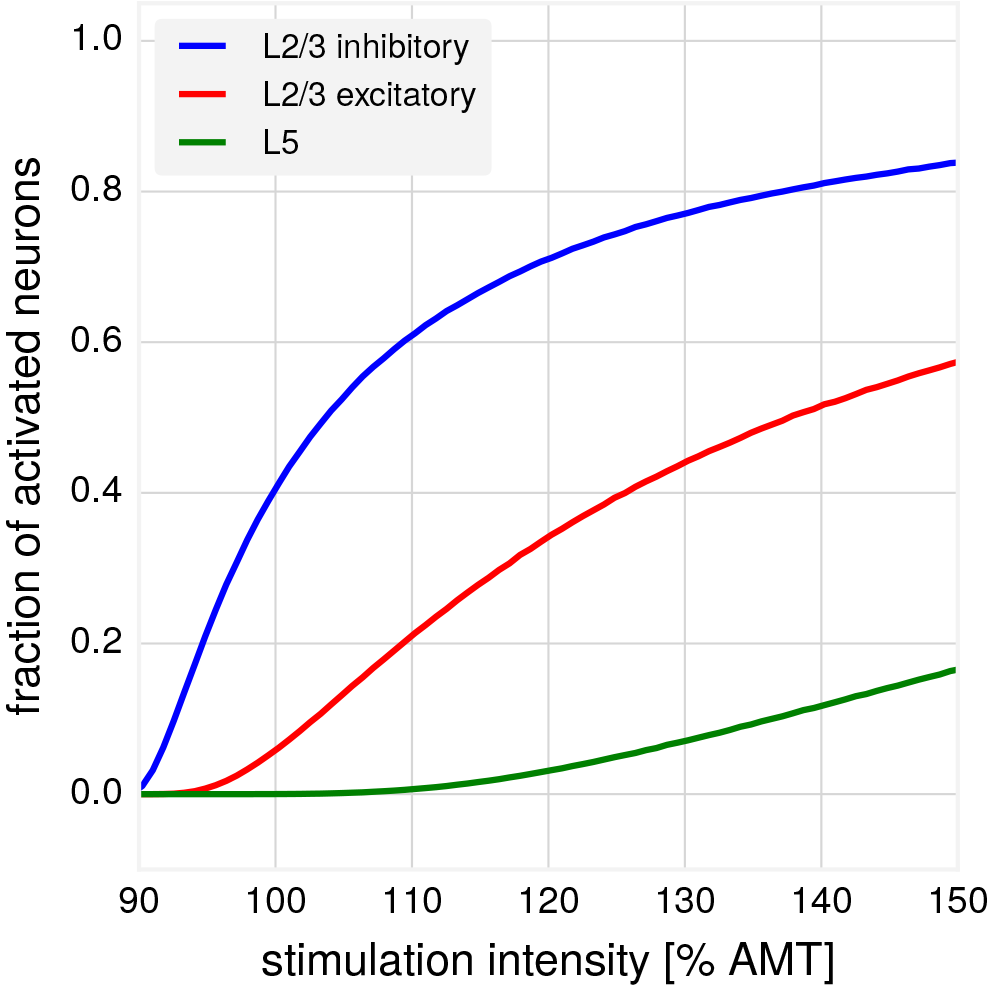
Proportion of directly activated neurons for different stimulation intensities as a percentage of the active motor threshold. Recruitment is stimulation-intensity dependent, with the implemented assumption that inhibitory L2/3 neurons are more easily recruited than L2/3 excitatory neurons. In comparison, L5 neurons have the lowest probability to be directly activated by TMS.

To enable comparisons to experimental recordings, γ is translated to a scale based on the active motor threshold (AMT). Experimental I-wave recordings performed at varying intensities [33, 32, 34] show that at AMT typically only a small *I*_1_-wave is visible, while the response tends to saturate at four I-waves for increasing intensity. Therefore, we chose two γ values to correspond to these situations, with linear interpolation in between. However, as recordings show large variability between subjects, as well as sensitivity to parameters such as location and orientation or type of the coil used, this mapping is to be viewed as a rough approximation.

### Model implementation

The model is implemented in Python and NEURON [35]. The time step for numerical integration is set to 0.025 ms. Each stimulation starts after a settling time of 300 ms during which the rhythmic input to the L2/3 neurons entrains oscillatory activity. Results are obtained as means of 25 simulations, with error bars showing the standard deviation. For additional details regarding the model structure and all parameter settings, see Table 1. More details concerning the model derivation can be found in [16].

## Results

### I-wave characteristics and their dependence on stimulation intensity

In a first step, we established that the model with spontaneous background activity is still able to produce the characteristic I-waves seen in epidural recordings. Examples of generated I-waves for different simulated stimulation intensities are compared to experimentally obtained data (replotted from [23]) in Fig. 4. Increasing the stimulation intensity increases the number of I-waves and their amplitude. Using a common multiplicative gain factor for recruitment of excitatory and inhibitory neurons produces a sigmoidal model response (see Fig. 5), as it becomes harder to recruit additional neurons for increasing stimulation intensities. Here we pooled over random phases, simulating TMS stimulation without EEG-feedback. Exemplarily, the high oscillatory power condition was used here. The scale parameter for the exponential distribution from which the current for injection is drawn was adjusted to match experimental results [23, 34]. The dependence on stimulation intensity is captured qualitatively by both, the spike measure as well as the I-wave amplitude measure.

**Figure 4:**
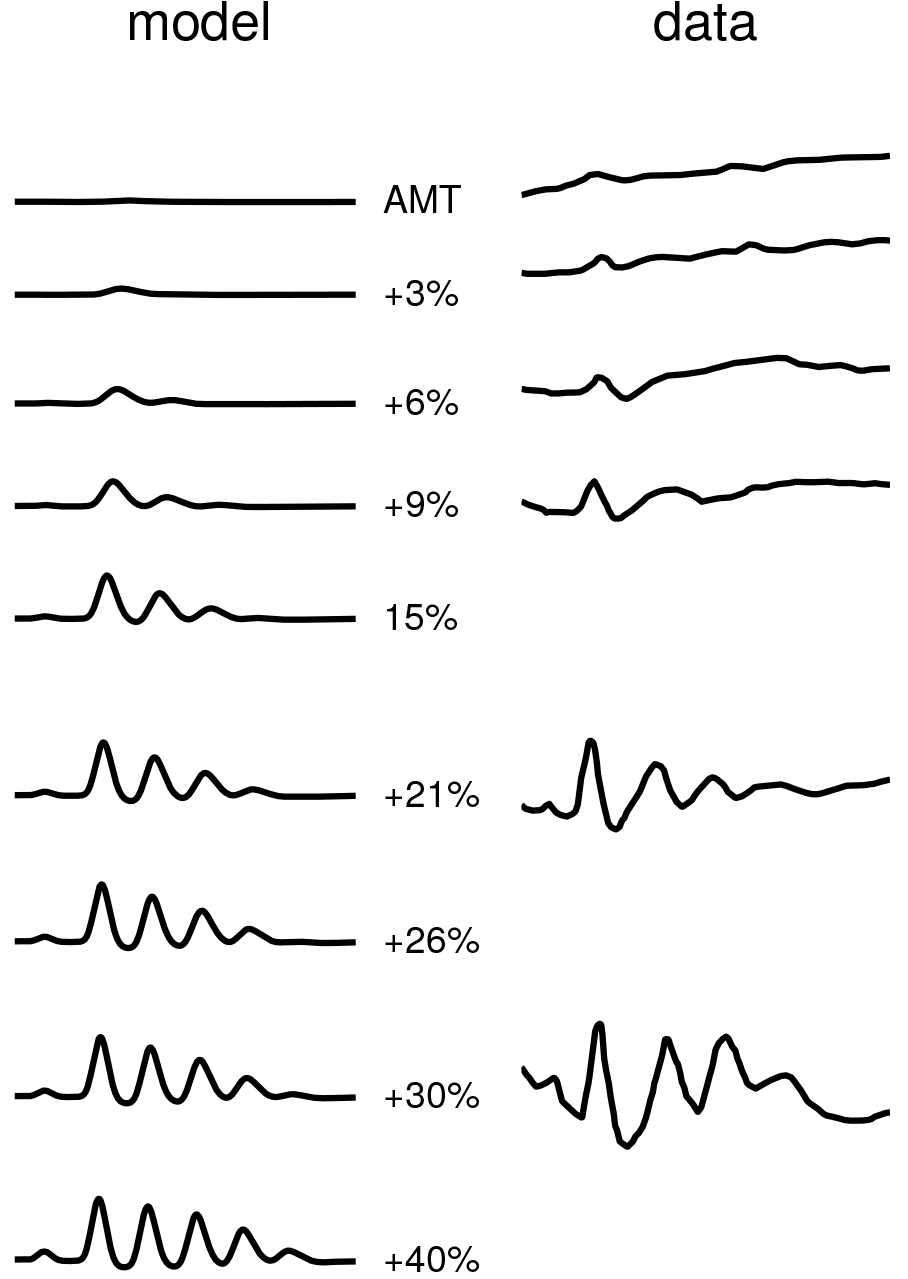
Left: Examples of I-waves generated by the model for different stimulation intensities. The number and amplitude of generated I-waves increases as a function of the stimulation intensities. Results shown are for 10 trials, pooled over random phases. Right: I-waves measured experimentally, data replotted from [23]. Averaged over 10 trials.

**Figure 5:**
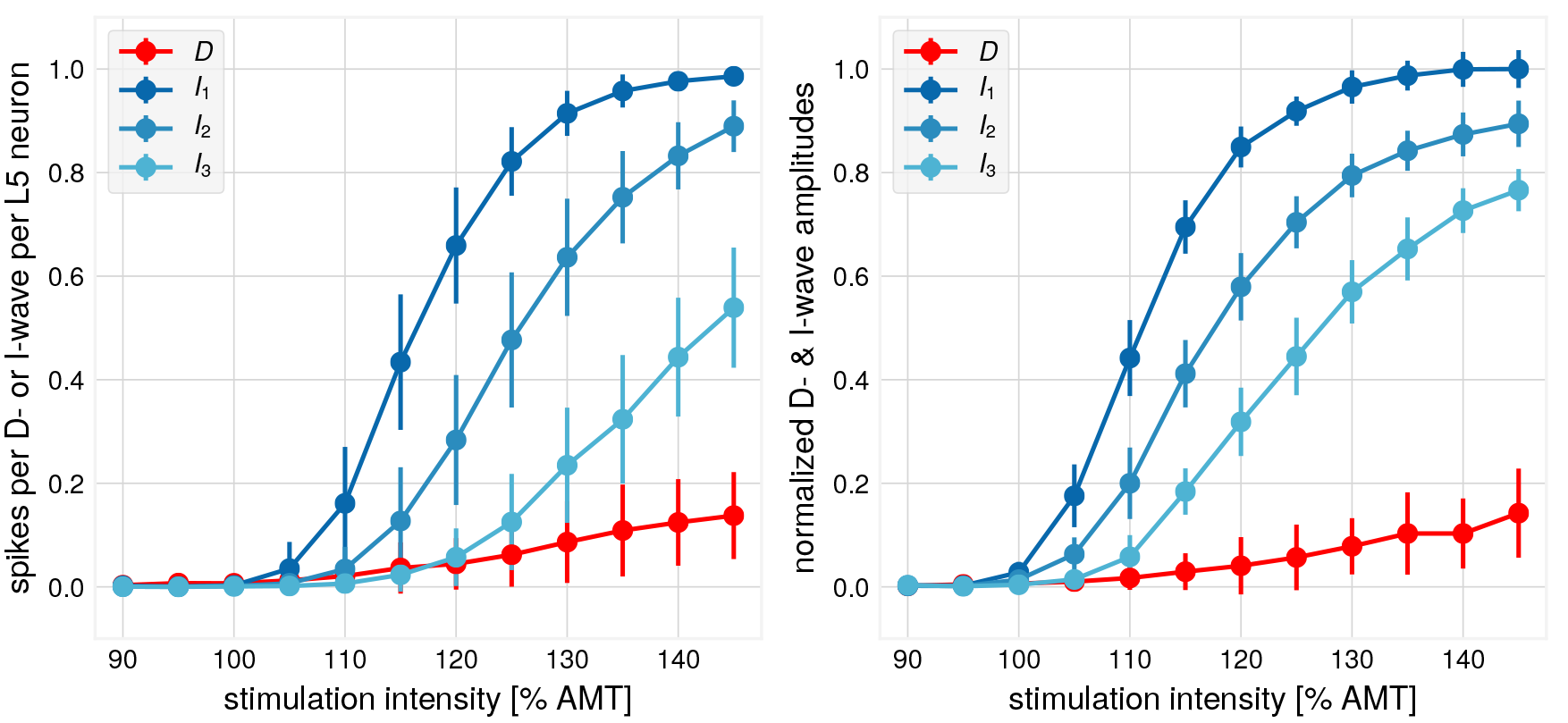
Input-output-curve. Left: number of spikes as a measure. Right: peak amplitude values as a measure. Amplitudes are normalized by the maximum mean value which was attained for the I_1_-wave. The number of spikes as well as peak amplitude values of generated I-waves increase with increasing stimulation intensity, with qualitatively similar behavior. Number of trials: 25, TMS onset at random phases, error bars represent one standard deviation.

### Modulation by oscillatory phase

In a next step, we investigated how the model’s response to TMS depends on the precise time of stimulation relative to the phase of the network’s oscillatory background activity. We first consider the case of high oscillatory power. We found that applying a TMS pulse at a specified phase of the background stimulation affects the number of generated spikes and the amplitude of I-waves differently. The model shows systematic modulation of the number of generated L5 spikes depending on the phase at which the TMS pulse is applied, as seen in Fig. 6 (Top). The largest number of spikes is elicited when the peak of L5 somatic subthreshold oscillations coincides with the timing of the TMS pulse (peak phase bin 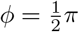). The difference between peak and trough response (phase bin with 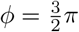) depends on stimulation intensity, with larger differences for lower stimulation intensities. The more background activity is present at the time of the TMS pulse, the more spikes can be elicited, as the TMS pulse is more likely to push neurons across their the action potential threshold towards spiking. L5 neurons which generate a large number of spikes in response to TMS are more likely to have a more elevated somatic membrane potential before stimulation onset in comparison to L5 neurons with a small number of TMS-induced spikes, see Fig. 7. The large standard deviation in this figure indicates that pre-TMS membrane potential is only one factor contributing the response to TMS in the model.

**Figure 6:**
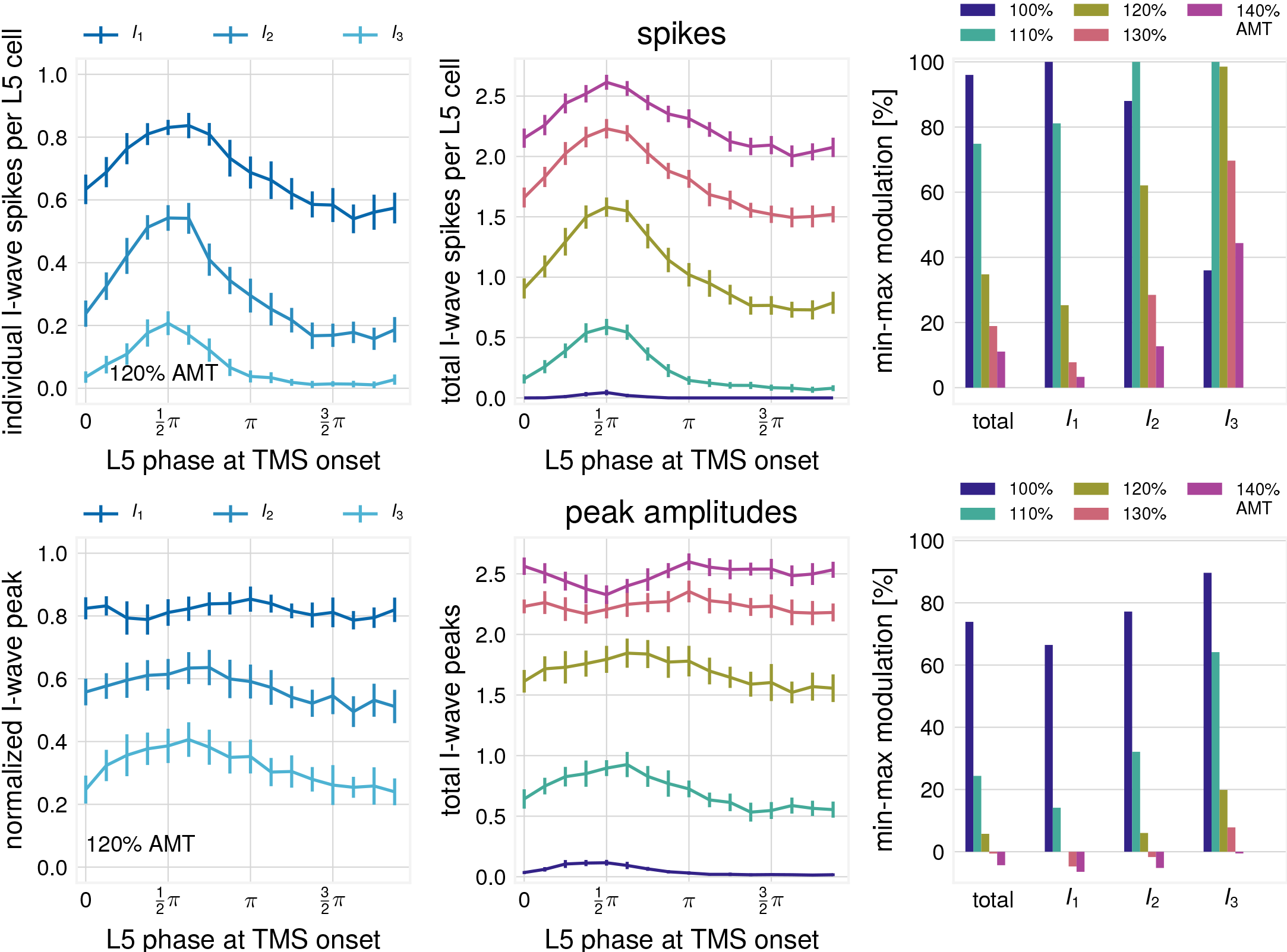
Phase-modulation as evaluated by different measures. Mean over 25 trials for each phase bin, error bars show one standard deviation. Top: Phase-modulation as evaluated by spike counts. Top left: absolute I-wave spike number is modulated by phase, shown for the first three I-waves. Top middle: sum of the first three I-wave spikes is modulated by phase and stimulation intensity. Top right: Phase-modulation of spikes constituting respective I-waves, as evaluated with 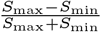. 100 with *S*_max_ as the number of spikes in phase bin 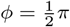 and *S*_min_ in phase bin 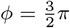. Bottom: Phase-modulation as evaluated by peak I-wave amplitude. Bottom Left: I-wave amplitude for the first three I-waves. Normalized by highest observed I-wave peak. Bottom Middle: sum of the first three I-wave amplitudes per stimulation intensity. Bottom Right: Phase-modulation, evaluated with peak values analogous to the top right figure.

**Figure 7:**
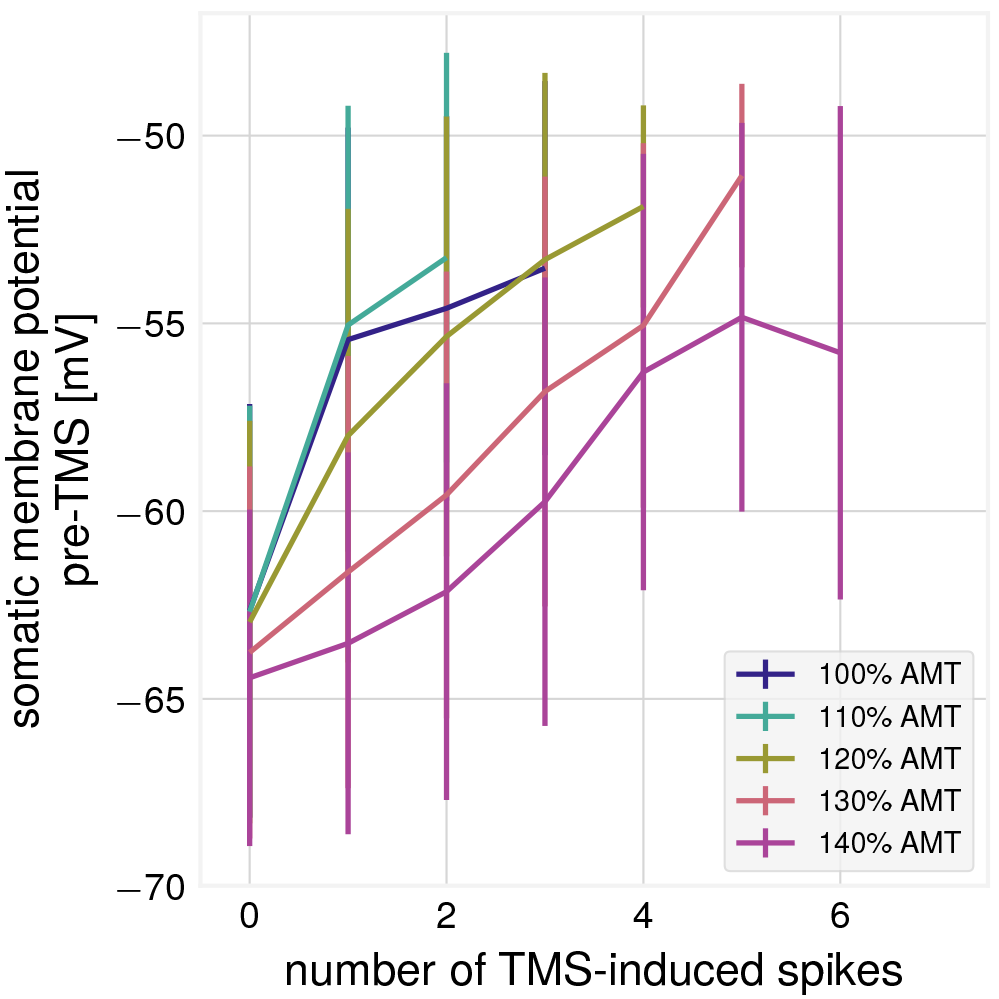
TMS-induced spikes plotted against pre-stimulation somatic membrane potential (mean over 5 ms before stimulation), for different intensities. Error bars show one standard deviation. All L5 neurons are pooled. Many spikes tend to be generated, when the membrane was strongly depolarized prior to the TMS pulse.

In contrast, Fig. 6 (Bottom) shows that I-wave amplitudes show comparatively little modulation by phase, depending on the stimulation intensity. For early I-waves, no clear differentiation between peak maxima and peak minima of L5 excitability can be seen for larger intensities. Only the I_3_-wave amplitude shows a clear amplitude modulation between peak and trough phase conditions in the expected direction when comparing to the spike measure. For 100% AMT, only in the peak phase condition a response will be elicited and no spikes in the trough phase condition, resulting in a large value of the min-max modulation index.

The number of generated spikes for peak and trough phase conditions is shown in Fig. 8 which illustrates how a smaller number of spikes can result in unchanged I-wave amplitude. A neuron is defined as active, if it fires in the respective I-wave interval. For the I_1_-wave, even though the overall number of active neurons is reduced by 9.1%, the number of spikes at the peak I_1_-time differs only by a small amount between conditions, which results in a similar I-wave peak amplitude. Only when spike timing relative to I-wave peak and number of spikes differ greatly, as for the I_3_-wave, the I-wave peak amplitude is changed between conditions. Correspondingly, a smaller number of spikes with less dispersion can yield larger I-wave amplitudes as compared to larger number of spikes with higher dispersion. This demonstrates a varying degree of sensitivity of the spikes and I-wave amplitude measures to changes in spontaneous activity.

**Figure 8:**
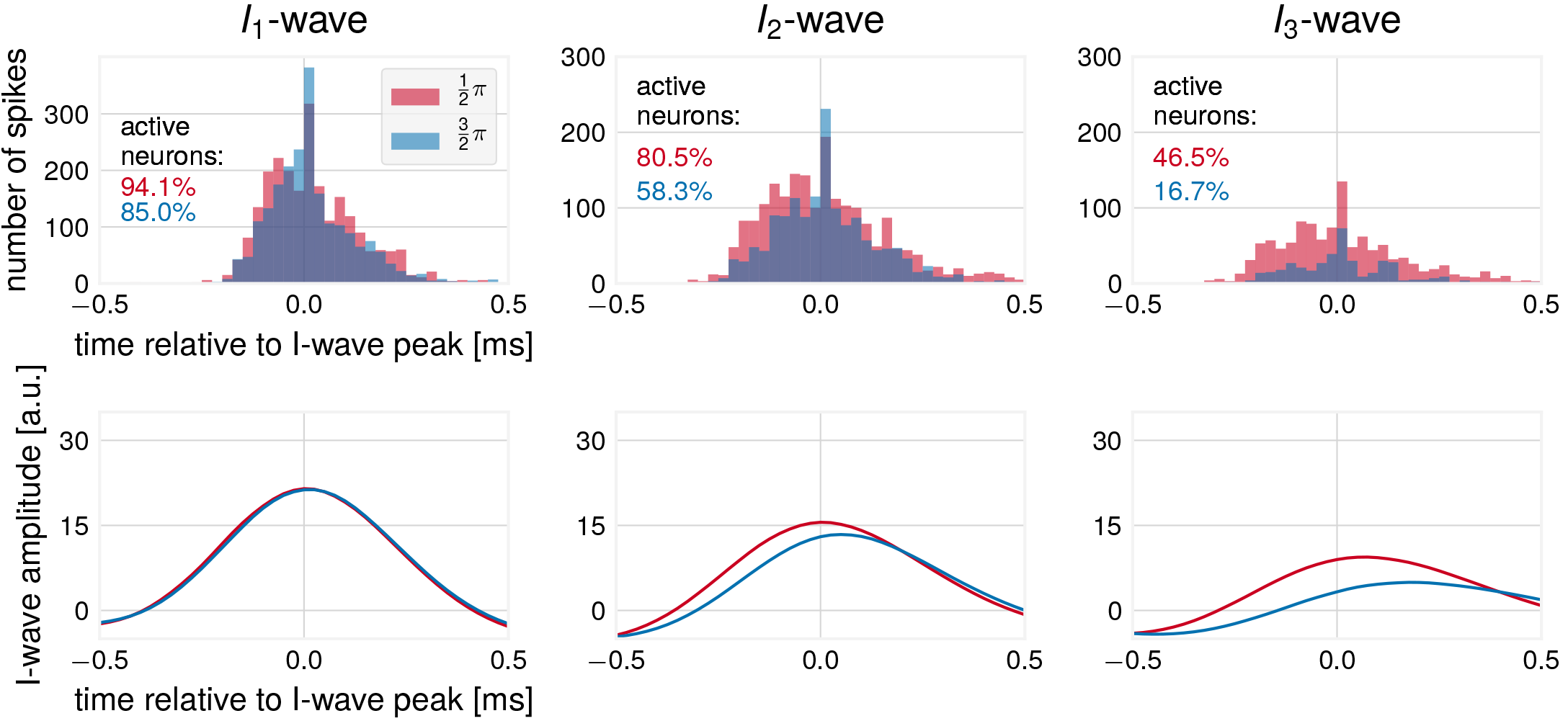
Illustration of difference in spike number and I-wave amplitude, for peak phase 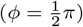 and trough phase 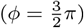 conditions. Top: I-wave spikes for individual I-waves. For this plot, all spikes from all L5 neurons were pooled. Bottom: I-wave time-courses. Only when the number and timing of spikes differs substantially, differences in I-wave amplitude can be seen.

### Modulation by oscillatory power

Finally, we tested for the effect of oscillatory power. The different conditions are illustrated in Fig 2. As shown in Fig. 9, if the oscillatory rhythm is pronounced with clearly differentiable high-activity and low-activity states, the response will be strongly modulated by the phase of the oscillatory rhythm. This results in an increased variability of the spike number elicited by a single pulse, when collapsing over phases.

**Figure 9:**
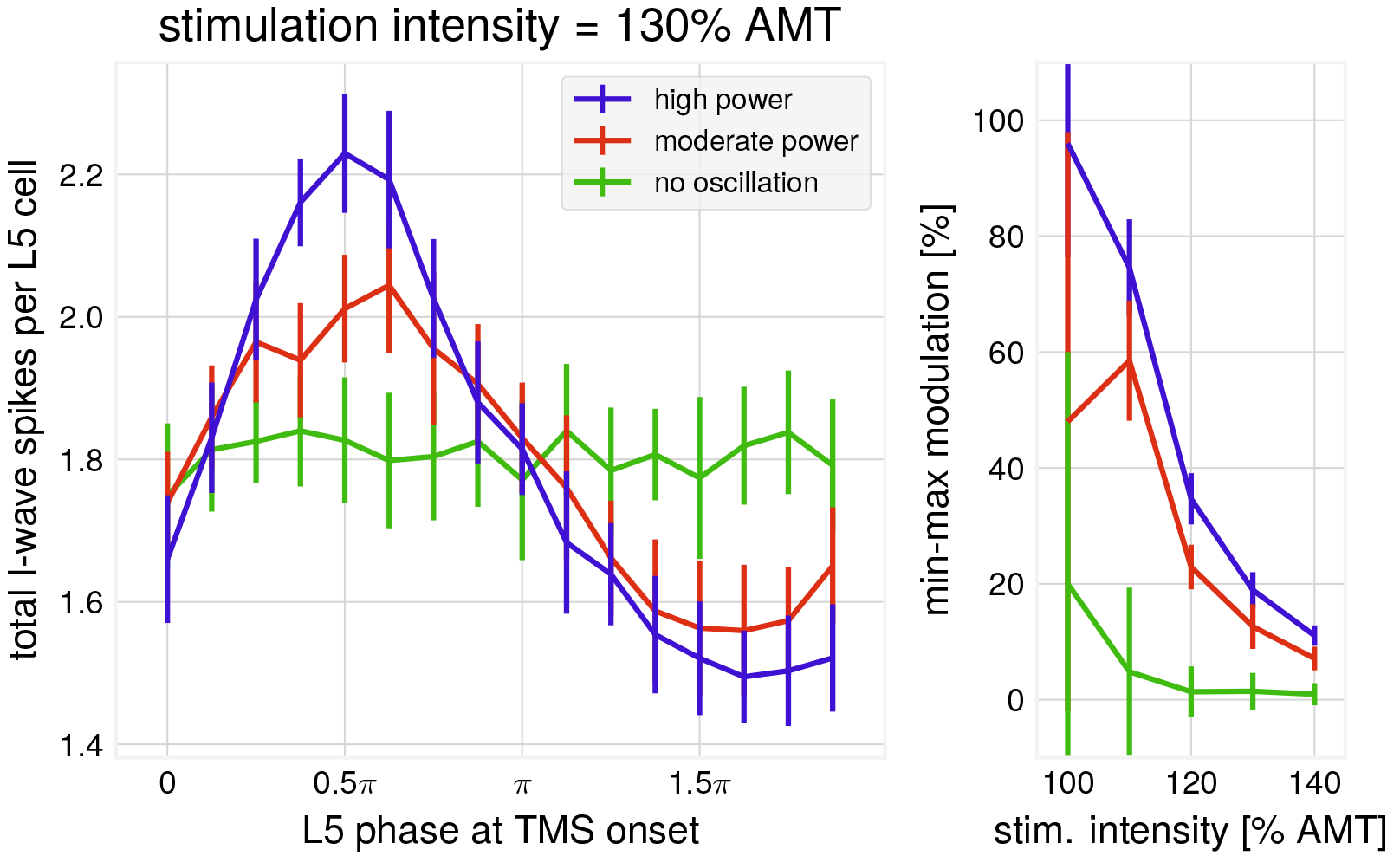
Phase-modulation for different oscillatory power conditions as evaluated by spike number. Left: Differences between power conditions for stimulation intensity = 130% AMT. The difference is more pronounced for higher oscillation power and leads to more variability in spike number for high oscillation power. Mean over 25 trials for each phase bin, error bars show one standard deviation. Right: Phase-modulation index (as in Fig. 6) for all intensities. The highest degree of modulation is achieved for high power of the ongoing rhythm and small stimulation intensities. Mean over 25 trials for each data point, error bars show one standard deviation.

## Discussion

Successful clinical application of TMS is hampered by the large and unexplained variability in the effects of various plasticity induction protocols. Since activity drives plasticity [36], much of this variability may be explained by factors inducing variability in cortical responses such as the brain state at the moment of stimulation. Our model reveals interactions between TMS responses and ongoing cortical oscillatory background activity and shows how cortical rhythms can contribute to the variability of TMS effects. The model is able to produce realistic I-waves and shows a dependence of TMS responses on power and phase of ongoing brain oscillations. It predicts that the strongest TMS response can be obtained for stimulation at the moment when L5 pyramidal neurons are maximally depolarized.

### Differentiation between numbers of spikes and I-wave amplitudes

Our results indicate that in the interplay with spontaneous activity, a differentiation has to be made between a spike measure and an I-wave measure of TMS effects. The number of generated L5 spikes shows a clear modulation according to the phase, while the corresponding effect on I-wave amplitudes can be smaller. We argue that this is related to a previously observed phenomenon: unchanged I-wave dynamics for vastly different MEP amplitudes. For certain TMS protocols [37, 38], large changes in MEP amplitude can be measured, but the corresponding I-waves are hardly changed. As shown in Fig. 8, different numbers of spikes can produce the same I-wave amplitudes. This suggests that I-waves are a less sensitive read-out measure compared to the total number of generated spikes. Looking at individual I-waves, the model suggests that later I-waves are more sensitive to changes in experimental conditions, as a large reduction of the number of spikes is needed to reduce I-wave amplitude significantly. But it is unclear how a high level of synchrony for later I-waves can be preserved with a reduced I_1_-wave. Interestingly, reliable modulation of early I-waves without changes in later I-waves was observed for the cTBS protocol [39]. Further investigation of experimental conditions where specifically the I_1_-wave is modulated would therefore be of interest. While the investigation of I-waves has given valuable insights into cortical circuits that contribute to the behavioral output [15], the interpretation remains difficult. The motor system is assumed to be based on rate coding [40, 41], so multi-unit recordings should be more predictive of MEP characteristics, compared to I-waves. However, there is recent evidence for an effect of precise spike timing on behavior [42]. As I-waves occur on a very fast time scale, simultaneous recordings of L5 spiking activity, I-waves, and muscle activity should reveal which measure is more predictive of MEP amplitude.

### Modulation by oscillation phase and relation to TMS-EEG

The model shows a clear dependence of TMS-response on pre-stimulation activity, with larger responses in a phase of larger L5 neuron depolarization. But there is no trivial correspondence between the phase of L5 membrane potential oscillations and the phase of the *μ*-rhythm, as measured by electroencephalography. The EEG signal is recorded with multiple electrodes on the scalp. In the sensor space, neighboring electrodes are necessarily related through volume conduction, but will exhibit phase-shifts [43]. The signal from a spatial filter constructed from sensor space may be shifted with respect to the phase of the generators of the *μ*-rhythm. The location of the functional areas generating the rhythm will vary between subjects [44], and also between frequency bands [45, 46]. Therefore, individualized filters may have to be used. Since the inverse problem of EEG has many solutions [47], attempts using source-reconstructed filters will have to make assumptions about the orientation of the dipole. This may also contribute to a mismatch between the phase of the targeted *μ*-generators and the global phase according to which the TMS stimulation is applied. Knowing that maximum MEP amplitude will be evoked at a phase where L5 neurons are maximally depolarized could help to identify the μ-generators and constrain the construction of effective spatial filters [48]. The dependence of phase-modulation on intensity could help to explain inter-individual differences, as experimentally, stimulation intensity is chosen with respect to subject individual criteria, such as resting or active motor threshold, size of MEPs, or subject comfort level. In general, the model predicts the largest modulation of TMS responses by oscillation phase when the stimulation intensity is low.

### Modulation by oscillation power and the role of subthreshold oscillations

The degree of phase-modulation is also dependent on the degree to which oscillatory rhythms are present in the motor system of the subject. The more pronounced the rhythms are, the more the TMS response is modulated by the phase of the ongoing activity. It is unclear, however, if different levels of EEG power in a frequency band for different subjects are due to less power in the neural activity, or a different orientation of the dipole making it harder to pick up the signal [49, 50]. This uncertainty about the true oscillatory power in the neural activity impedes inter-subject comparisons. In addition, the individual location of the corresponding motor areas has to be considered. Furthermore, membrane oscillations are modulated by task-related activity [30]. Overall, we predict less variability of MEPs at lower power of the generators of the oscillations.

### Model limitations

Our model has a simple feed-forward structure. While this appears sufficient for modeling the responses to single-pulse TMS, the addition of recurrent connections may be required for capturing how reverberating activity due to early TMS pulses influencing the responses to later pulses. This is an interesting topic for future work.

## Acknowledgements

The authors acknowledge support from the Quandt foundation.

## Methods Summary

Here we provide a summary of the neural network model in table form, following the guidelines proposed in [51]. For a full specification, see the available model code at http://TBD.

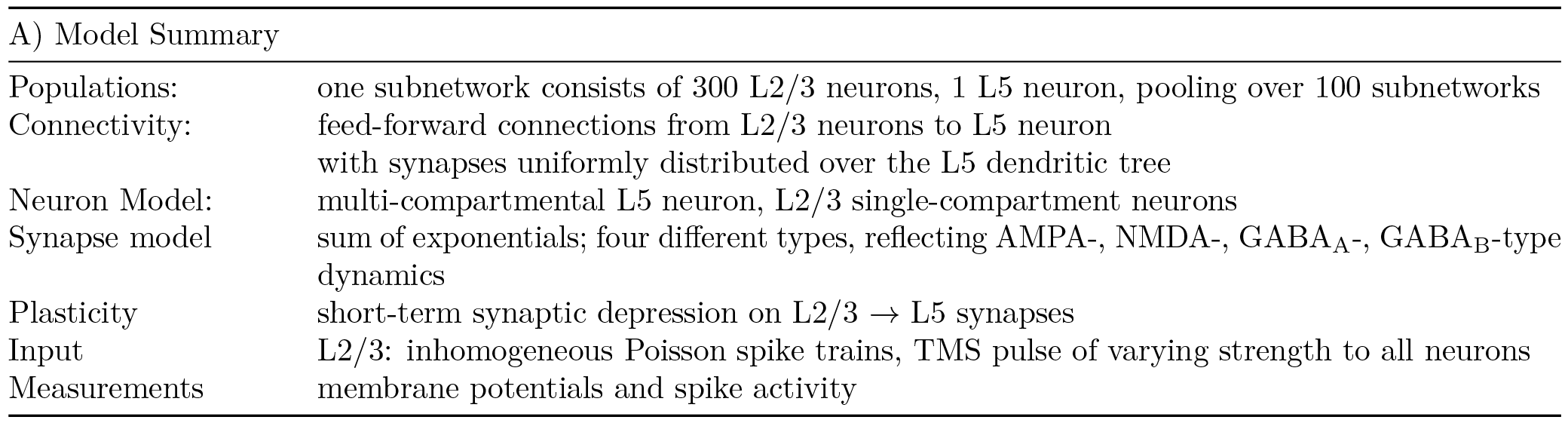

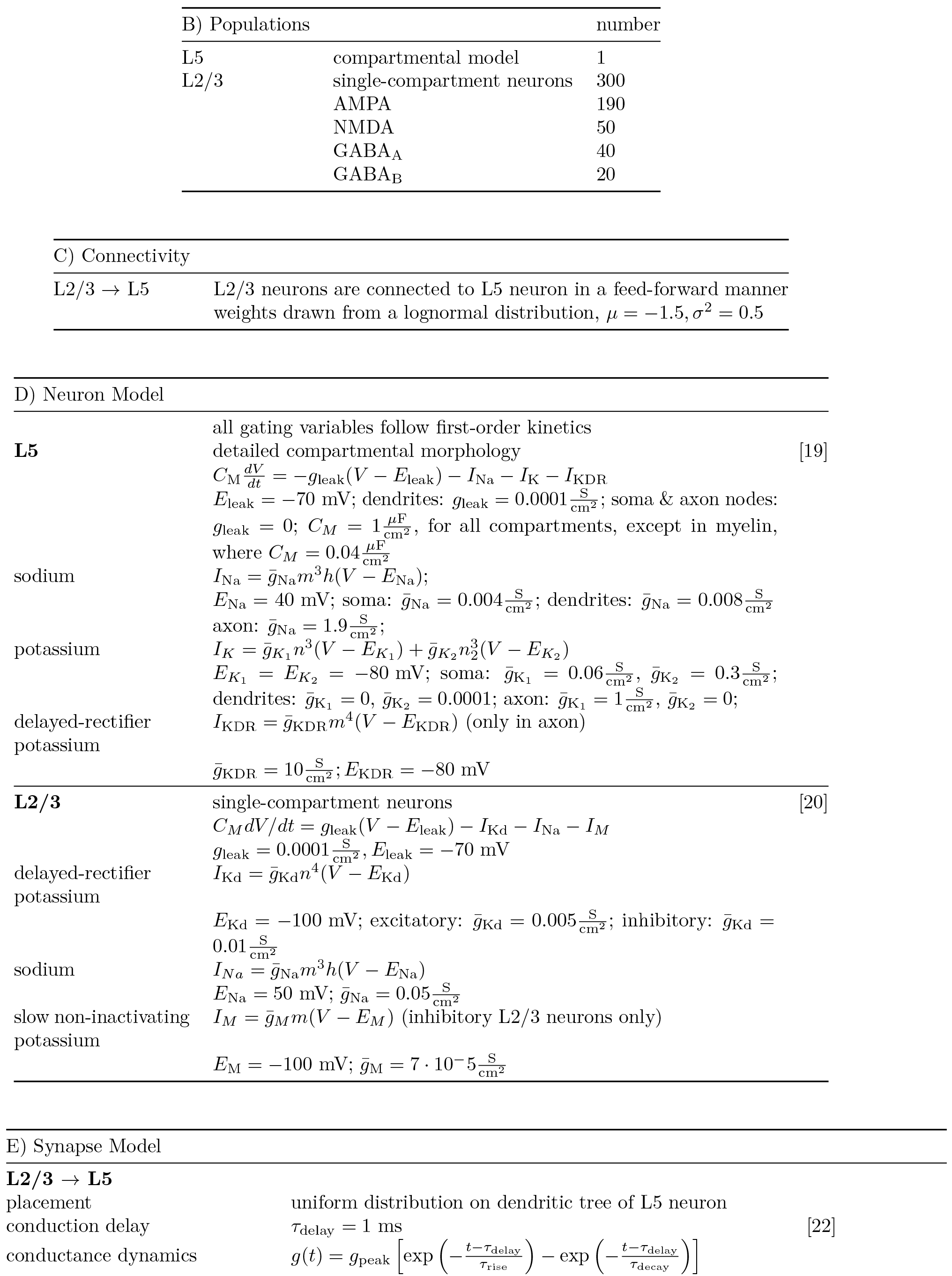

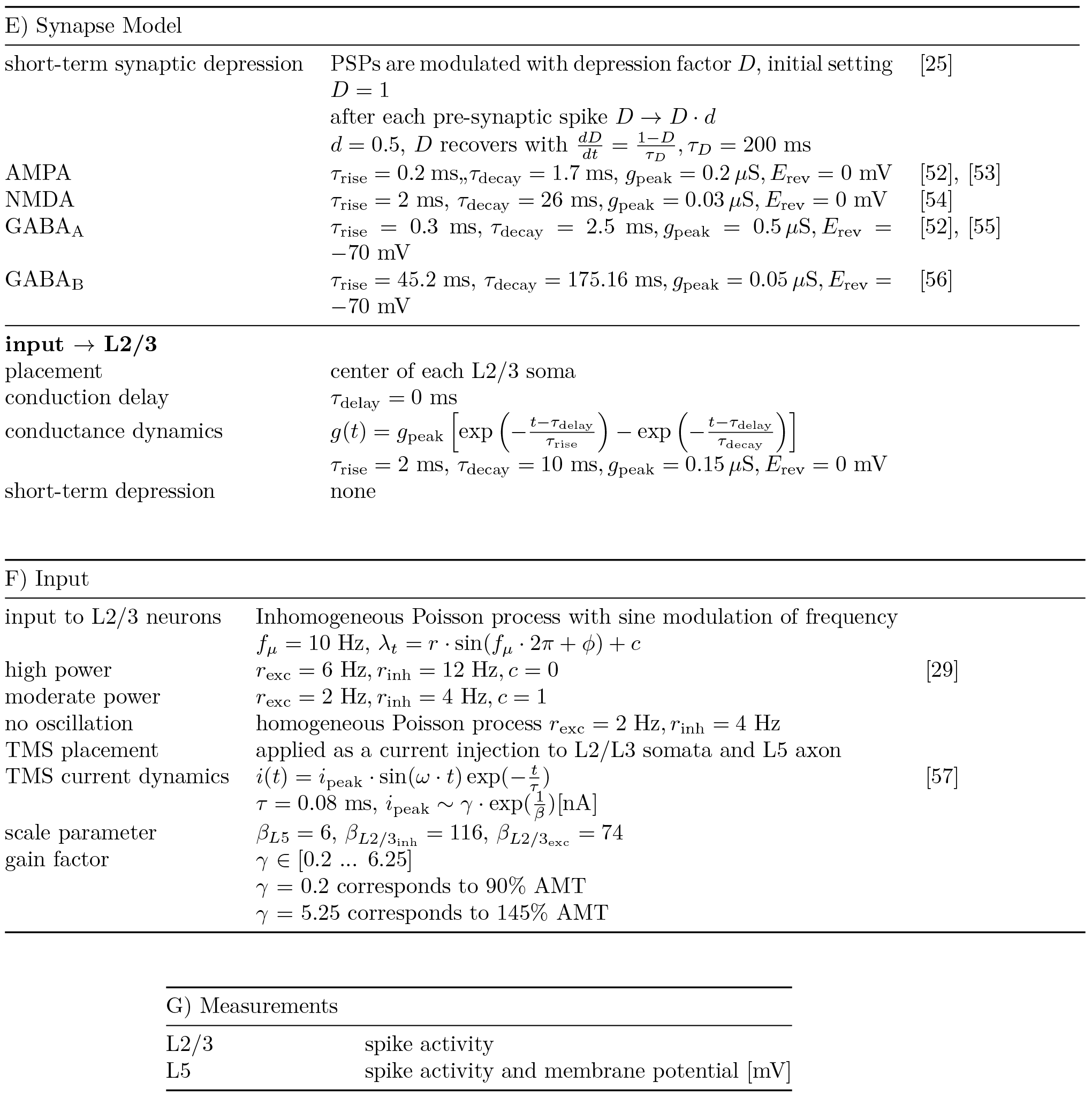

